# Oxygen kinetics during CXL using symmetrically and asymmetrically pulsed UV-irradiation

**DOI:** 10.1101/2022.08.17.504256

**Authors:** Maria A. Komninou, Malavika H. Nambiar, Beatrice E. Frueh, Volker Enzmann, Philippe Büchler, Theo G. Seiler

## Abstract

**Purpose:** To investigate oxygen kinetics during symmetrically pulsed and asymmetrically pulsed crosslinking (p-CXL) with and without supplementary oxygen at different irradiances and corneal depths.

**Design:** Experimental, laboratory study

**Methods:** In de-epithelialized porcine eyes, a femtosecond-laser generated tunnel was used to place a fibre-probe in corneal depths of 200 and 300 µm to measure the local oxygen concentration. After riboflavin imbibition, the corneas were irradiated at 9, 18 and 30 mW/cm^2^ for 10 seconds On and 10 seconds Off; while the oxygen concentration was continuously measured until oxygen levels depleted below the oxygen sensor’s threshold (1%) or until stabilized. All experiments were performed under normoxic (21%) and hyperoxic (>95%) conditions and the obtained data were used to identify parameters of a numerical algorithm for oxygen consumption and diffusion. Following the algorithm’s development, the suggested asymmetrical pulsing values were experimentally tested. For 9, 18 and 30 mW/cm^2^ the suggested tested pulsing schemes were 3 seconds On : 9 seconds Off, 2 seconds On : 9 seconds Off and 1 second On : 9 seconds Off respectively.

**Results:** The minimum, available stromal oxygen for p-CXL in normoxic environment was decreasing <1% for 9, 18 and 30 mW/cm^2^ in 200 and 300 μm. Using optimized p-CXL, the minimum available oxygen increased to 3.8, 1.8 and 2.8 % at 200 μm, for irradiances of 9, 18 and 30 mW/cm^2^, respectively, where the periods exhibited an equilibrium state. At 300 μm, 1.1 % of oxygen was available for 30 mW/cm^2^. Using a hyperoxic environment, the oxygen concentration was 19.2% using 9 mW/cm^2^ in 200 μm, dropping to 17.0% in 300 μm. At 18 mW/cm^2^, the concentrations were 3.9% and 1% in 200 and 300 μm, respectively. Using 30 mW/cm^2^, all oxygen was depleted below the threshold limit (1% O_2_) for both depths. Using optimized pulsing in combination with hyperoxic environment, the oxygen concentration was 42.0% using 9 mW/cm^2^ in 200 μm and 43.3% in 300 μm. At 18 mW/cm^2^, the concentrations were 24.7% and 16.1% in 200 and 300 μm, respectively. Using 30 mW/cm^2^, the minimum oxygen availability was 25.7% and 13.7% in 200 and 300 μm, respectively.

**Conclusion:** Supplementary oxygen during symmetrical and asymmetrical p-CXL increased the oxygen availability during corneal cross-linking. The pulsed irradiance and the hyperoxic environment potentially increased the efficacy of corneal cross-linking in deeper corneal layers and higher irradiances. The numerical algorithm for asymmetrical pulsing led to the quantification of “On” and “Off” times related to different scenarios such as irradiances.

## Introduction

Corneal crosslinking (CXL) is a safe and effective technique to prevent progression of keratectasia such as keratoconus^1,2^ or post-LASIK ectasia with low rates of complication^3^. Reactive oxygen species (ROS) are produced when the photosensitizer riboflavin is activated by UV irradiation, forming new bond in the cornea’s extracellular matrix^4^. Conventional epithelium-off crosslinking has already demonstrated the safety and long term-efficacy needed for stabilization of progressive keratectasia^5–8^. However, it has been shown that in deeper stromal layers of the cornea, all available O_2_ is depleted within seconds and the rediffusion is not sufficient to achieve effective aerobic CXL^9^ under normoxic conditions. During the last years, pulsed CXL (p-CXL) has been proposed as an alternative to continuous UV-irradiation considering that the on-off pattern will allow more oxygen to rediffuse into the corneal stroma and will lead to an enhanced delivery of oxygen allowing a more efficient and shorter treatment^10,11^. The theory of acceleration derives from the Bunsen-Roscoe law of reciprocity indicating that the same amount of photochemical reaction can be achieved by increasing UV-A irradiation and reducing time^8^. However, whether pulsed UVA light is more effective than continuous irradiation has not been widely examined and, therefore, remains questionable. Mazzotta et al. reported that the effect of pulsed CXL was superior to continuous CXL, contrariwise Toker et al. reported that there was no statistically significant difference between the 2 groups^12,13^.

The aim of this study was to investigate the oxygen availability, consumption and rediffusion during pulsed CXL under hyperoxic (>95% O_2_) and normoxic (21% O_2,_ room air) oxygen conditions at various depths using different irradiances and pulsing schemes, as well as to test an optimized pulsing scheme utilizing a predictive algorithm that could possibly enhance the clinical treatment.

## Materials and Methods

### Experimental Setup

The process for sample preparation, oxygen environment and cross-linking has been previously described in detail^9^. For this study, the porcine eyes were divided into following four groups (n = 6 for each group):

1. Symmetrical (10”:10”) pulsed CXL under normoxic conditions – The UV-irradiation was set to 10 seconds On:10 seconds Off
2. Asymmetrical pulsed CXL in normoxic conditions – The laser was configured to the optimized pulsed protocol (3”:9” for 9 mW/cm^2^, 2”:9” for 18 mW/cm^2^ and 1”:9” for 30 mW/cm^2^)
3. Symmetrical (10”:10”) pulsed CXL in hyperoxic conditions – The laser was set to 10 seconds On:10 seconds Off
4. Asymmetrical pulsed CXL under hyperoxic conditions– The laser was configured to the optimized pulsed protocol (3”:9” for 9 mW/cm^2^, 2”:9” for 18 mW/cm^2^ and 1”:9” for 30 mW/cm^2^)

The groups were subjected to CXL in an atmospheric and hyperoxic environment in corneal depths of 200 and 300 μm (Table 1). For more accurate data, the optic oxygen micro-sensor (NTH-PSt1; PreSens, Regensburg, Germany), connected to a microfiber optic oxygen meter (Micro TX3; PreSens) measured one data point per second. All investigated groups received epithelial removal, followed by the application of 0.1% riboflavin (Vibex Rapid, Avedro, Waltham, MA, USA) every 2 minutes for a total of 10 minutes and irradiation. Thereby, the UV-source (PXL Platinum 330, PeschkeTrade, Huenenberg, Switzerland) was focused onto the surface of the cornea delivering irradiances of 9, 18 and 30 mW/cm^2^. For both sets of experiments, the UV-irradiance was delivered in a pulsed form and the oxygen measurements were recorded until the oscillation period was constant.

**Table 1:** Experimental parameters applied for each test group

| Irradiance<br>(mW/cm<sup>2</sup>) | Depth<br>(μm) | p-CXL<br>10":10" |  |  | p-CXL asymmetrical |  |  | Continuous |  |  |
| --- | --- | --- | --- | --- | --- | --- | --- | --- | --- | --- |
|  |  | Time<br>(min) | min O<sub>2</sub> % |  | Time<br>(min) | min O<sub>2</sub> % |  | Time<br>(min) | min O<sub>2</sub> % |  |
|  |  |  | Hyperoxic | Normoxic |  | Hyperoxic | Normoxic |  | Hyperoxic | Normoxic |
| **9** | 200 | 20 | 19.2 | < 1 | 40 | 42.0 | 3.8 | 10 | 2.6 | < 1 |
|  | 300 |  | 17.0 | < 1 |  | 43.3 | < 1 |  | 1.2 | < 1 |
| **18** | 200 | 10 | 3.9 | < 1 | 27.5 | 24.7 | 1.8 | 5 | < 1 | < 1 |
|  | 300 |  | 1.0 | < 1 |  | 16.1 | < 1 |  | < 1 | < 1 |
| **30** | 200 | 6 | < 1 | < 1 | 30 | 25.7 | 2.8 | 3 | < 1 | < 1 |
|  | 300 |  | < 1 | < 1 |  | 13.7 | 1.1 |  | < 1 | < 1 |

### Statistical analysis

The figures creation, boxplot representation and average minimum and maximum values of the data were calculated using the Python programming language and the libraries matplotlib and plotly. Statistical analysis of the results was performed using a one-way ANOVA including Bonferroni’s post-hoc analysis (IBM SPSS Statistics Corp., Armonk, NY, USA). The p-values less than 0.05 were considered to indicate a statistical significance.

### Numerical Setup

Based on experimental data previously collected with continuous laser irradiations^9^, a pulsatile algorithm of O_2_ diffusion in the cornea was developed. The temporal profile of oxygen depletion during CXL treatment, at different intensities and the profile of oxygen increase, due to rediffusion when the laser was turned off, were used to estimate the oxygen levels at 200 µm and 300 µm for different pulsation strategies. This was done by interpolating the previously recorded oxygen concentration of the constant irradiance with the corresponding oxygen concentration from the “on” and “off” times of the pulsating laser. This algorithm was verified with pulsed CXL data for 10-second-intervals from the “On” and “Off” states. This algorithm was then used to determine the optimal pulse duration by testing different “On” and “Off” states. Since oxygen consumption is directly related to the number of cross-links generated per unit time, the optimal pulse strategy was defined to maximize the oxygen consumption predicted by the algorithm during the laser. The optimization was performed for 200 um and 300 μm depth for 9, 18, and 30 mW/cm^2^ irradiations, respectively. The algorithm was analyzed in 1D using MATLAB®.

## Results

### Validation of the numerical algorithm against standard pulsed CXL data

The pulsation algorithm was validated using experimental data with the symmetrical 10”:10” p-CXL protocol in normoxic conditions. After oxygen had dropped below the sensor’s threshold (Table 2), the numerical predictions were within the standard deviation of the oxygen measurements for the irradiances of 9, 18, and 30 mW/cm^2^ at both 200 µm and 300 µm depths (Figure 1). Therefore, the algorithm was then used to determine the pulsation strategy that maximized the oxygen consumption rate, which yielded an optimum of 3 seconds On:9 seconds Off at 9 mW/cm^2^, 2 seconds On:9 seconds Off at 18 mW/cm^2^, and 1 second On:9 seconds Off at 30 mW/cm^2^.

**Table 2:** Irradiances, depths, time and protocols for which the O2 percentage drops below the sensor’s threshold limit.

| Irradiance<br>(mW/cm<sup>2</sup>) | Depth<br>(μm) | p-CXL<br>10":10" |  | p-CXL optimal |  |
| --- | --- | --- | --- | --- | --- |
|  |  | Below threshold depletion<br>time (sec) |  | Below threshold depletion<br>time (sec) |  |
|  |  | Hyperoxic | Normoxic | Hyperoxic | Normoxic |
| **9** | 200 | - | 8 | - | - |
|  | 300 | - | 6 | - | 4 |
| **18** | 200 | - | 4 | - | - |
|  | 300 | - | 3 | - | 3 |
| **30** | 200 | 6 | 3 | - | - |
|  | 300 | 8 | 2 | - | - |

**Table 3:** Boxplots that represent significant difference (<0.05) between each other+

| Boxplot name & number | Normoxic environment |  | Hyperoxic environment |  |
| --- | --- | --- | --- | --- |
|  | Comparison of boxplots for min values | Comparison of boxplots for max values | Comparison of boxplots for min values | Comparison of boxplots for max values |
| 1: 9 mW/cm<sup>2</sup> at 200 μm (10":10") | 1 – 7<br>1 – 8<br>1 – 9 | 1 – 5<br>1 – 6<br>1 – 10<br>1 – 11<br>1 – 12 | 1 – 2<br>1 – 3<br>1 – 5<br>1 – 6<br>1 – 7<br>1 – 10 | 1 – 3<br>1 – 6<br>1 – 10 |
| 2: 18 mW/cm<sup>2</sup> at 200 μm (10":10") | 2 – 7<br>2 – 8<br>2 – 9 | 2 – 5<br>2 – 6<br>2 – 11 | 2 – 4<br>2 – 7<br>2 – 8<br>2 – 9<br>2 – 10<br>2 – 11 | 2 – 7<br>2 – 10 |
| 3: 30 mW/cm<sup>2</sup> at 200 μm (10":10") | 3 – 7<br>3 – 8<br>3 – 9 | 3 – 7<br>3 – 8<br>3 – 9 | 3 – 4<br>3 – 7<br>3 – 8<br>3 – 9<br>3 – 10<br>3 – 11<br>3 – 12 | 3 – 4<br>3 – 7<br>3 – 8<br>3 – 9<br>3 – 10 |
| 4: 9 mW/cm<sup>2</sup> at 300 μm (10":10") | 4 – 7<br>4 – 8<br>4 – 9 | 4 – 7<br>4 – 8<br>4 – 9 | 4 – 5<br>4 – 6<br>4 – 7<br>4 – 10 | 4 – 5<br>4 – 6<br>4 – 10 |
| 5: 18 mW/cm<sup>2</sup> at 300 μm (10":10") | 5 – 7<br>5 – 8<br>5 – 9 | 5 – 7<br>5 – 8<br>5 – 9 | 5 – 7<br>5 – 8<br>5 – 9<br>5 – 10<br>5 – 11<br>5 – 12 | 5 – 7<br>5 – 8<br>5 – 9<br>5 – 10 |
| 6: 30 mW/cm<sup>2</sup> at 300 μm (10":10") | 6 – 7<br>6 – 8<br>6 – 9 | 6 – 7<br>6 – 8<br>6 – 9 | 6 – 7<br>6 – 8<br>6 – 9<br>6 – 10<br>6 – 11<br>6 – 12 | 6 – 7<br>6 – 8<br>6 – 9<br>6 – 10 |
| 7: 9 mW/cm<sup>2</sup> at 200 μm (optimized) | 7 – 8<br>7 – 9<br>7 – 10<br>7 – 11<br>7 – 12 | 7 – 10<br>7 – 11<br>7 – 12 | 7 – 8<br>7 – 9<br>7 – 10<br>7 – 11<br>7 – 12 | 7 – 11<br>7 – 12 |
| 8: 18 mW/cm<sup>2</sup> at 200 μm (optimized) | 8 – 9<br>8 – 11 | 8 – 10<br>8 – 11<br>8 – 12 | 8 – 9<br>8 – 12 | 8 – 12 |
| 9: 30 mW/cm<sup>2</sup> at 200 μm (optimized) | 9 – 10<br>9 – 11<br>9 – 12 | 9 – 10<br>9 – 11<br>9 – 12 | 9 – 10<br>9 – 12 | 9 – 12 |
| 10: 9 mW/cm<sup>2</sup> at 300 μm (optimized) | n.s. | n.s. | 10 – 11<br>10 – 12 | 10 – 11<br>10 – 12 |
| 11: 18 mW/cm<sup>2</sup> at 300 μm (optimized) | n.s. | n.s. | n.s. | n.s. |
| 12: 30 mW/cm<sup>2</sup> at 300 μm (optimized) | n.s. | n.s. | n.s. | n.s. |

**Figure 1:**
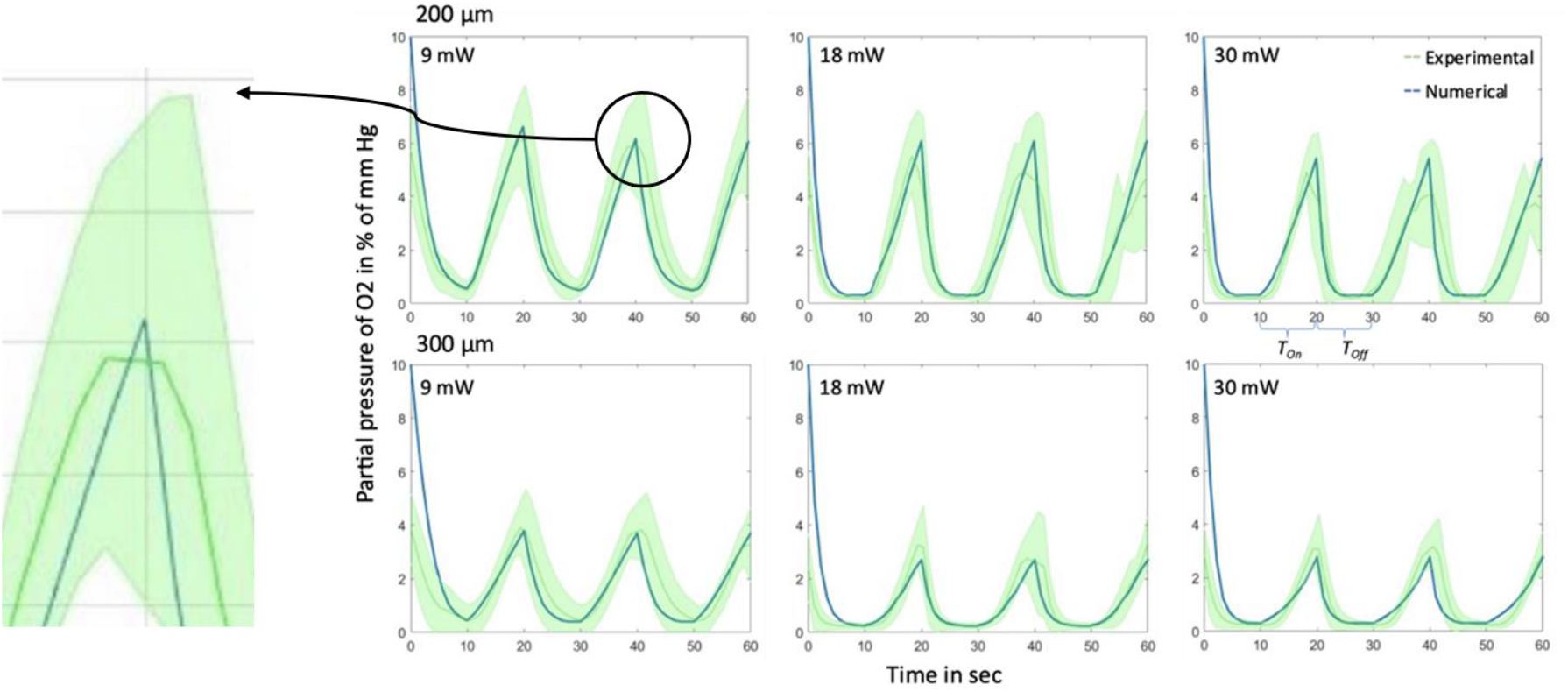
Validation of numerical algorithm against symmetrical (10”:10”) pulsed CXL experiments

### Oxygen concentrations during pulsed CXL treatment in normoxic and hyperoxic environments

Intrastromal oxygen measurements in a normoxic and hyperoxic environment were performed and are shown in Figure 2 and Figure 3, respectively. Thereby, blue curves show the oxygen consumption during cross-linking with an irradiance of 9 mW/cm^2^, the pink curves are of irradiance 18 mW/cm^2^ and the green curves represent the 30 mW/cm^2^ UV-irradiances. For both figures, subplots A and B represent the O_2_ consumption and rediffusion during the pulsing scheme of 10”:10” for 200 μm and 300 μm respectively, while subplots C and D represent the O_2_ consumption and rediffusion for the asymmetrical pulsing scheme.

**Figure 2:**
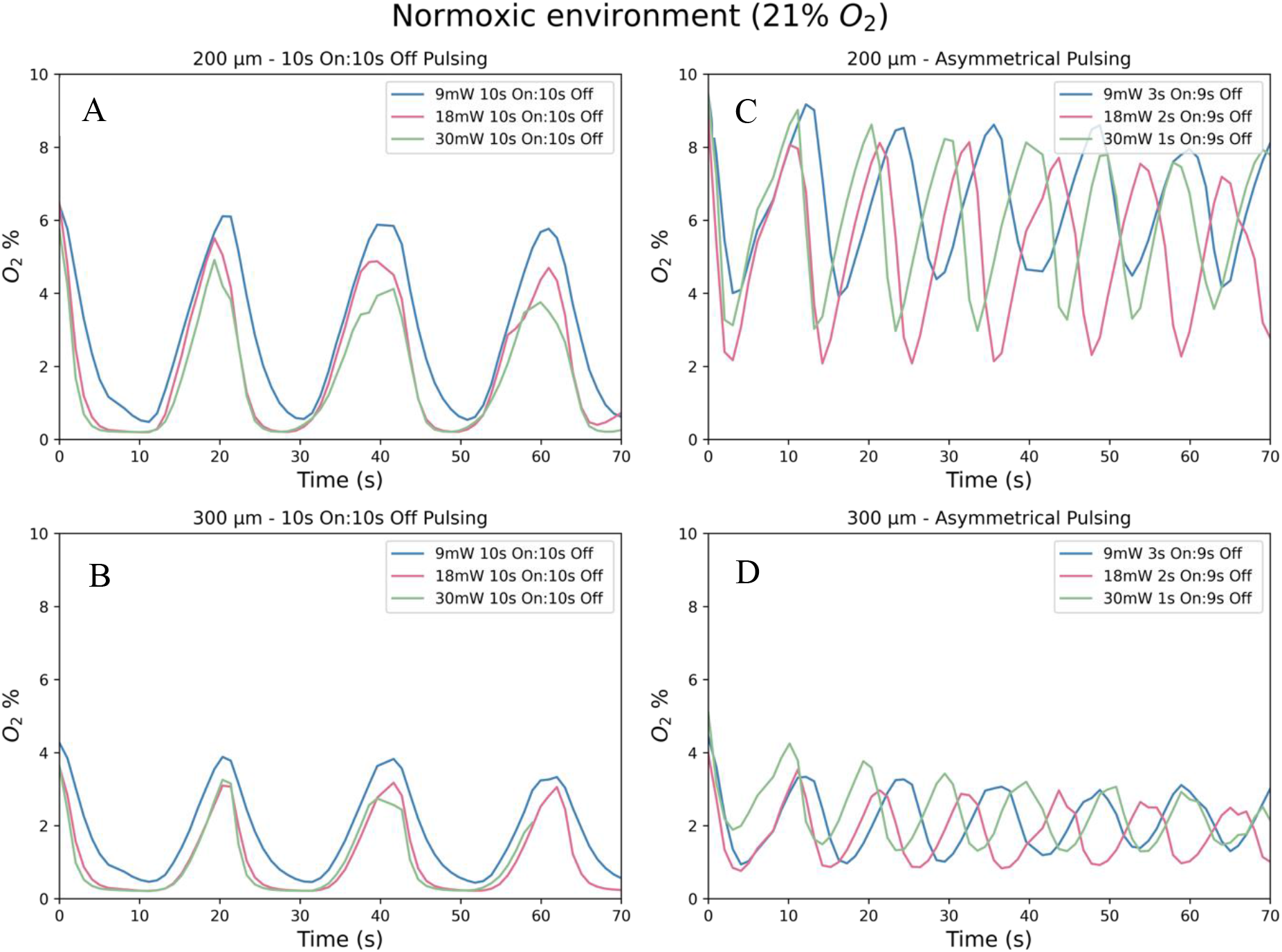
Average O2 measurements for normoxic environment. Panels A and B represent the 10’’:10’’ pulsing schemes whereas C and D represent the proposed asymmetrical pulsing scheme for 200 μm and 300 μm respectively

**Figure 3:**
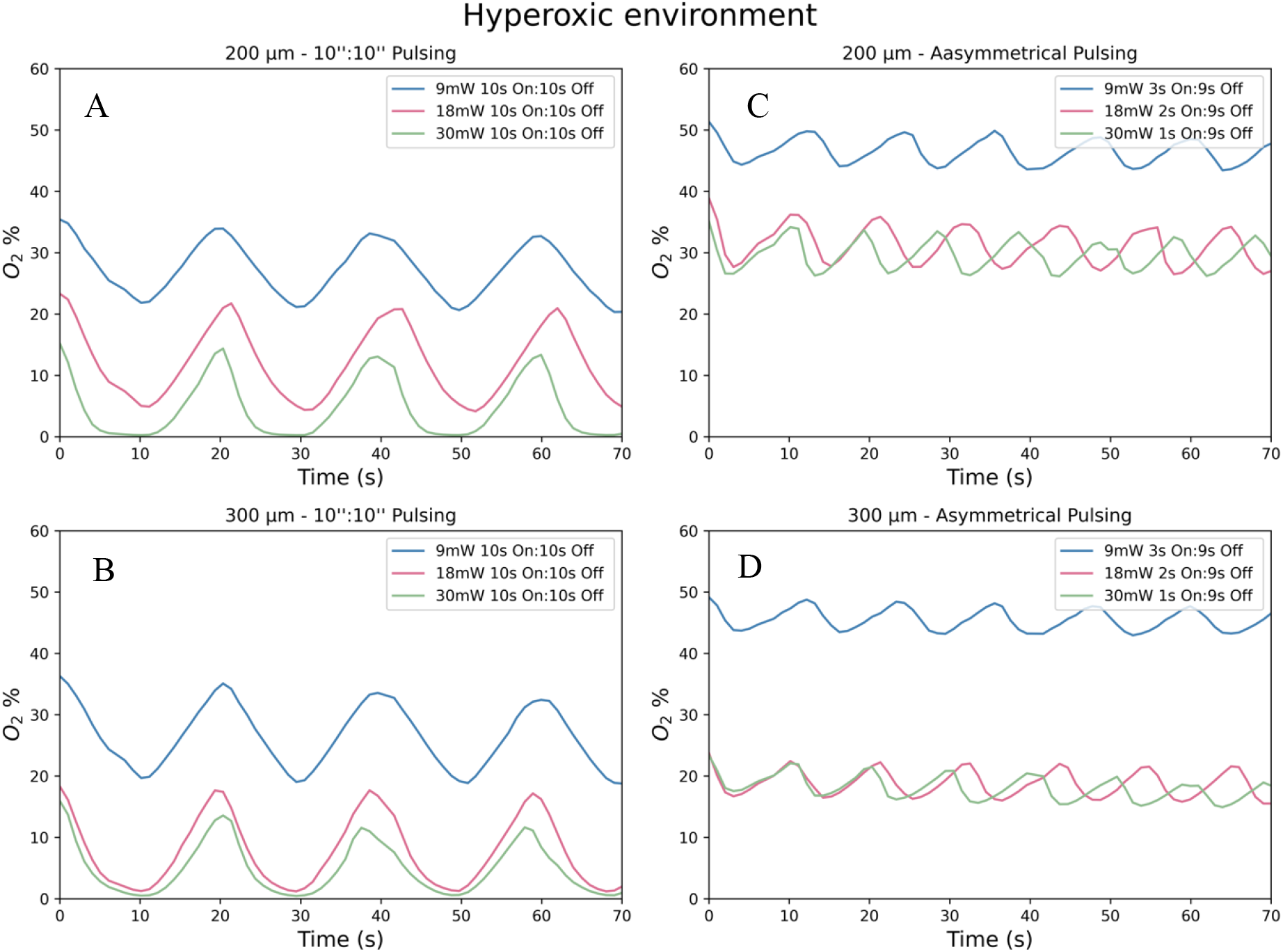
Average O2 measurements for hyperoxic environment. Subfigures A and B represent the 10’’:10’’ pulsing schemes whereas C and D represent the proposed asymmetrical pulsing scheme for 200 μm and 300 μm respectively

Once the UV irradiation was initiated, oxygen levels quickly descended. Table 1 shows the minimum values of the pulsing periods in percentage of oxygen and the total treatment time for the symmetrical and asymmetrical pulsing schemes, for hyperoxic and atmospheric oxygen environments. For the normoxic environment, the highest oxygen availability was found in the 9 mW/cm2 subgroups. In 200 μm depth the minimum oxygen was 3.8% when the asymmetrical pulsing scheme was used (Figure 2 C). For higher irradiances and deeper corneal layers the oxygen levels were below the sensor’s threshold. The statistical analysis for 200 μm showed statistical significance between minimum oxygen values for 10”:10” pulsing and optimized pulsing (p <0.001 between same irradiances) while for 300 μm there was no statistical significance between the two pulsing schemes.

Hyperoxic environment significantly increased the stromal oxygen availability (Figure 3). In 200 μm depth the minimum oxygen was 42.0% when the asymmetrical pulsing scheme was used and 19.2% for the symmetrical pulsing. For higher irradiances and deeper corneal layers, decreasing oxygen levels were observed. During the UV-off time of the periods, oxygen levels were rediffusing to the initial levels. The post-hoc analysis between the two pulsing groups, in 200 μm showed p < 0.001 between same irradiance groups. The same p-value was found for the 300 μm, when comparing the different pulsing schemes between same irradiances (9 and 18 mW/cm^2^) and for 30 mW/cm^2^ the p-value was equal to 0.002.

### Minimum and maximum values for normoxic and hyperoxic environment

Based on the minimum and maximum values of the periods, boxplots for 10’’:10’’ symmetrical pulsing and asymmetrical pulsing for 200 μm and 300 μm in normoxic and hyperoxic environment, respectively, were created (Figure 4 and Figure 5). Dark blue boxplots represented the highest values and light blue boxplots represented the lowest values of the periodic graphs for 200 μm. Dark red boxplots represented the highest values and light red boxplots represented the lowest values of the periodic graphs for 300 μm. For the 200 μm depth of the left figure the same six eyes had been tested. Similarly, another set of six eyes had been tested for 300 μm depth of the left figure. The same process was followed for the optimized pulsing scheme on the right figure.

**Figure 4:**
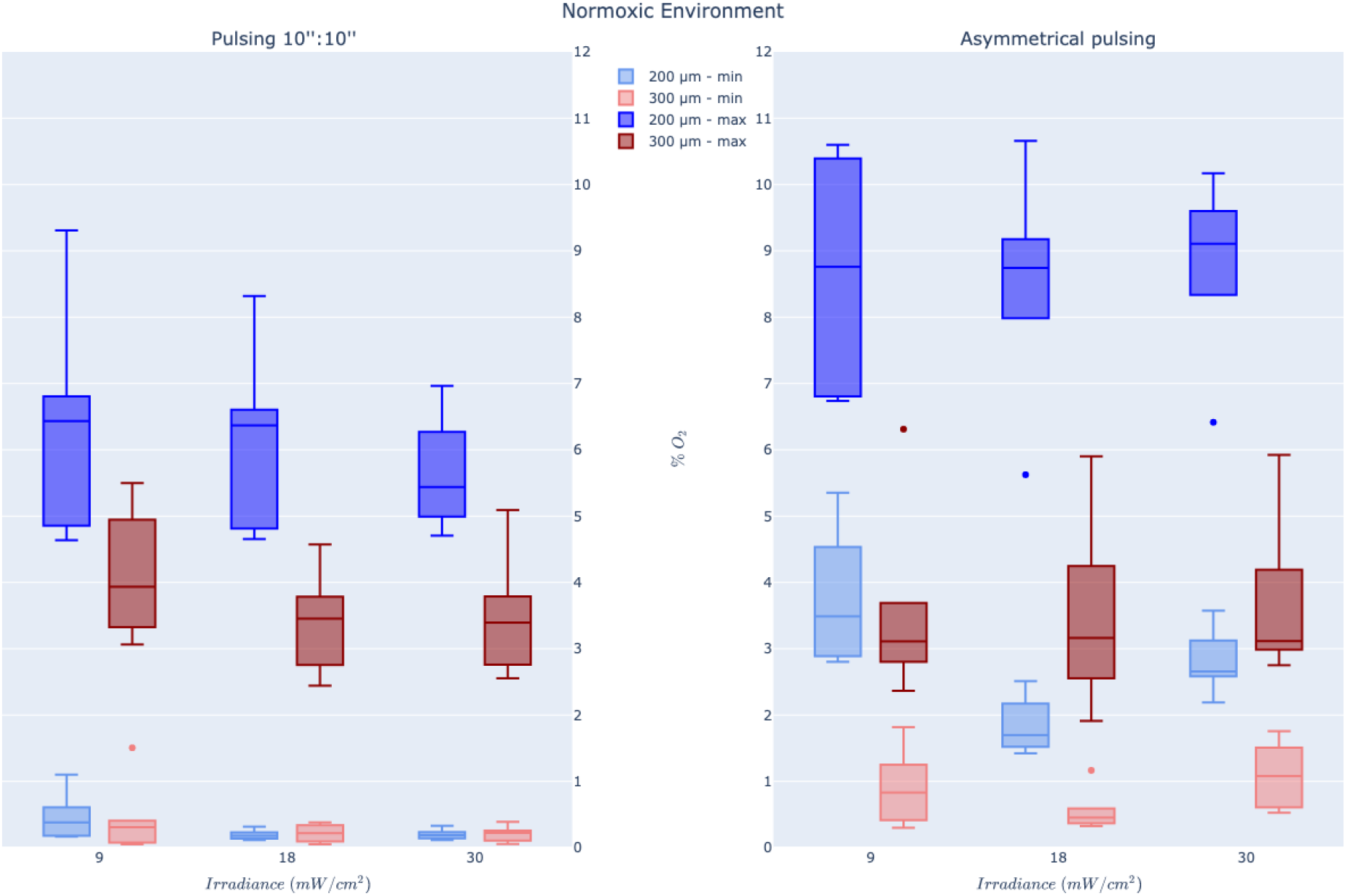
Boxplots for 10’’:10’’ pulsing (left) and asymmetrical pulsing (right) in normoxic environment for 200 μm and 300 μm. Dark blue boxplots represent the highest values and light blue boxplots represent the lowest values of the periodic graphs for 200 μm

**Figure 5:**
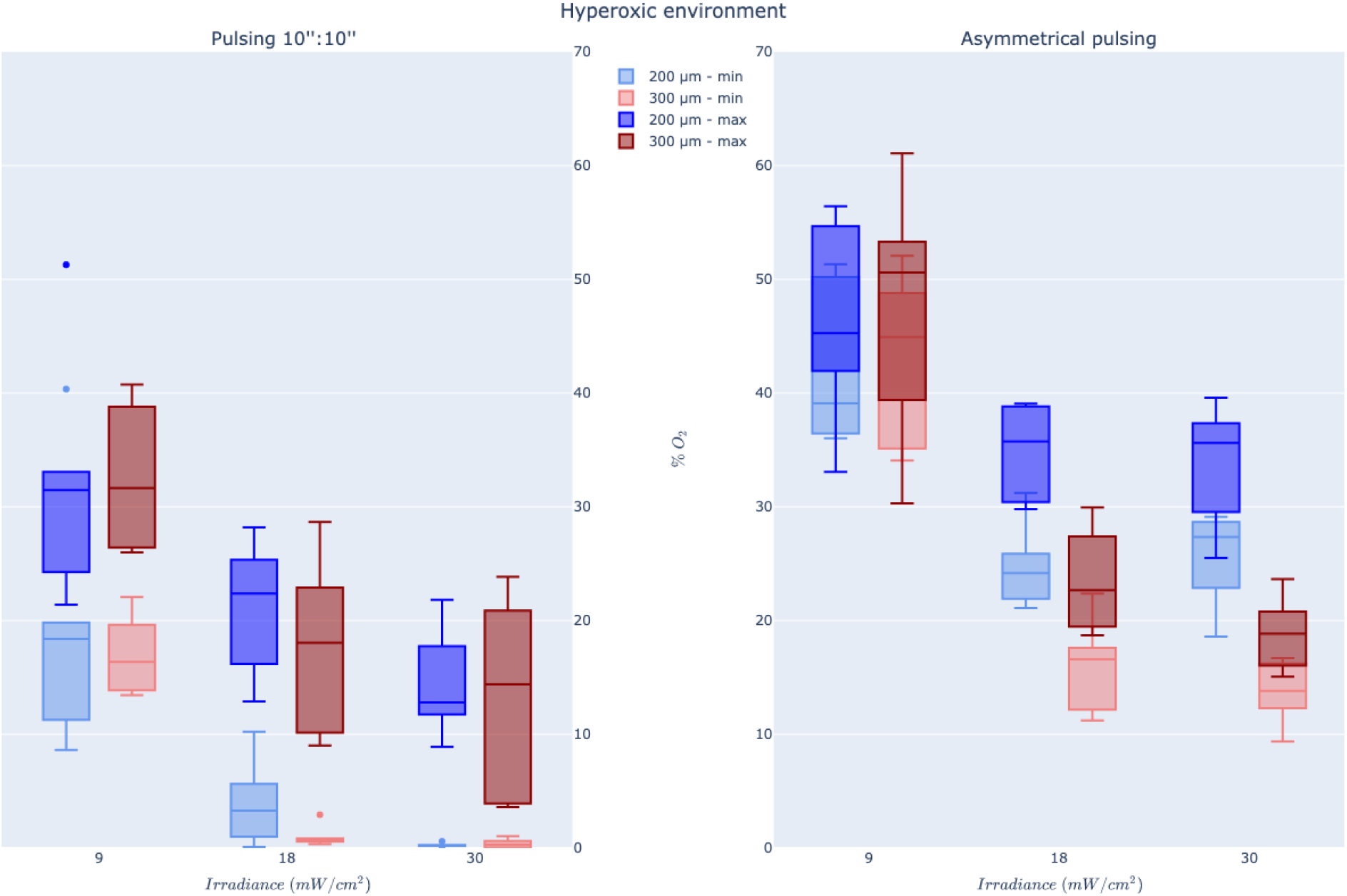
Boxplots for 10’’:10’’ pulsing (left) and asymmetrical pulsing (right) in hyperoxic environment for 200 μm and 300 μm. Dark blue boxplots represent the highest values and light blue boxplots represent the lowest values of the periodic graphs for 200 μm

For Figure 4 and the minimum values of the graphs’ periods during 10”:10” pulsing, the boxplots had almost the same median but different distributions. The boxplot for 9 mW/cm^2^ at 200 μm had a wider third quartile, while the rest five were relatively short suggesting a high level of similarity between each other. All six of them were well below the sensor limit of one percent. The minimum values of the graphs’ periods during asymmetrical pulsing showed an increase of the median for both 200 and 300 μm. The oxygen levels during asymmetrical pulsing were below the threshold limit for 9 and 18 mW/cm^2^, whereas for 30 mW/cm^2^ it was 1.1% at 300 μm and therewith above it.

Supplementary oxygen during treatment is increasing the available stroma oxygen, something that is shown in Figure 5.

For the minimum values of the graphs’ periods during 10”:10” pulsing, the medians for 9 mW/cm^2^ were 18.4 and 16.3 for 200 and 300 μm, respectively. These values are much higher than the normoxic environment for the same pulsing scheme. For 18 mW/cm^2^ at 300 μm depth and 30 mW/cm^2^ for both depths, the boxplots are not widely spread with the values agreeing to each other and the medians are below one, indicating that there is not enough oxygen for cross-linking. On the other hand, for the asymmetrical pulsing scheme, for 9 mW/cm^2^ at 200 μm the median is located at 38% O_2_ while for the 300 μm it is located at 45%.

### Experimental parameters for all experimentally tested groups

Table 1 represents a summary of all experimental parameters, like irradiance, corneal depth, treatment time and the resulting minimum oxygen concentrations during the periods of treatment but are also compared to the continuous data that had been acquired from our previous work^9^. The minimum values of oxygen derive from computing the minimum values of each eye separately and the averaging them.

Oxygen dynamics are distinctly different when conducted under hyperoxic conditions. During normoxic treatment at 200 μm depth with the 10”:10” pulsing scheme, the oxygen depletes below the sensor’s measuring limit in 8 seconds, 4 seconds and 3 seconds for 9 mW/cm^2^, 18 mW/cm^2^ and 30 mW/cm^2^, respectively. For 300 μm the O_2_ depletes even faster. During hyperoxic treatment, the oxygen depletes below the sensor limit only for the irradiance of 30 mW/cm^2^, for both depths. For 200 μm the O_2_ depletes within 6 seconds and for 300 μm the O_2_ depletes in 8 seconds (Table 2). Interestingly, for the asymmetrical pulsing scheme the oxygen depletes below the threshold only for 9 and 18 mW/cm^2^ at 300 μm depth (Table 1).

## Discussion

Oxygen is of great importance for the efficiency of CXL treatment. Indeed, previous studies have shown that performing CXL in a hyperoxic environment leads to an increase of available stroma oxygen and therefore a deeper and more effective treatment^9,14^. Simultaneously, previous studies have shown that p-CXL with a fluence of 7.2 J/cm^2^ leads to a deeper apoptotic effect^13,15,16^. To our knowledge, the oxygen consumption during 10’’:10’’ and asymmetrical p-CXL with a fluence of 5.4 J/cm^2^, in normoxic and hyperoxic environments has never been examined before. The results presented here demonstrate that a stable hyperoxic environment during symmetrical but especially asymmetrical pulsed treatments can counteract challenges associated with maintaining aerobic conditions as oxygen is faster consumed in accelerated protocols. Even in corneal depths of 200 and 300 μm, or higher irradiances like 9 – 30 mW/cm^2^ the available oxygen is adequate for treatment.

The data for 10’’:10’’ p-CXL in normoxic environment demonstrate that, regardless of irradiance and depth, performing CXL in a symmetrical pulsed form with an energy of 5.4 J/cm^2^ causes total depletion of stromal oxygen (Figure 2 A-B). During the rediffusion time in 200 μm, the O_2_ increases up to 6.41 %, 6.18 % and 5.63 % for 9, 18 and 30 mW/cm^2^. While, in 300 μm, the O_2_ increases up to 4.12 %, 3.41 % and 3.5% for the same irradiances. Given the treatment time that is needed for reaching the energy of 5.4 J/cm^2^ this pulsing protocol does not provide adequate oxygen for fast and efficient treatment since the time needed is double comparing to normoxic A-CXL time (Table 1) and for both protocols the oxygen drops below the oxygen threshold. During the asymmetrical pulsing scheme (Figure 2 C) the availability of oxygen for 200 μm corneal depth increases. During depletion time, the oxygen concentration does not decrease below 3.76 % for 9 mW/cm^2^, 1.84 % for 18 mW/cm^2^ and 2.8 % for 30 mW/cm^2^. During rediffusion, the concentration reaches the maximum value of the period in 8.5 seconds, for all irradiances. On the other hand, for 300 μm the lowest values of oxygen during depletion are below the threshold limit (1%) showing no improvement comparing it to the 10’’:10’’ p-CXL (Figure 2 D).

The use of hyperoxic environment for 10’’:10’’ and asymmetrical p-CXL, significantly improves the availability of oxygen even in higher irradiances and deeper corneal layers, something that is not feasible with continuous irradiations. For the symmetrical, 10’’:10’’ scheme in 200 μm, the minimum oxygen concentration lays at 19.2 % for 9 mW/cm2, at 3.9 % for 18 mW/cm2 and below 1 % for 30 mW/cm2 (Figure 3 A-B)

This corresponds to a 40-fold increase for 9 mW/cm^2^ and a 20-fold increase for 18 mW/cm^2^ when compared to the same depth and pulsing scheme, for normoxic environment. Simultaneously, the rediffusion concentration exhibits a 5-fold increase for 9 mW/cm^2^, a 3.4 fold increase for 18 mW/cm^2^ and 2.5 fold increase for 30 mW/cm^2^ comparing to the same depth and pulsing scheme, for normoxic environment. For 300 μm, the minimum oxygen concentration lays at 16.97 % for 9 mW/cm^2^ but for 18 mW/cm^2^ and for 30 mW/cm^2^ the amount of oxygen is well below the sensor’s threshold. This shows that even hyperoxic environment is not adequate for CXL in layers deeper than 300 μm and irradiances higher than 9 mW/cm^2^.

The asymmetrical pulsing scheme in hyperoxic environment (Figure 3 C-D) increases significantly the availability of oxygen. During oxygen depletion for 200 μm corneal depth, the concentration does not drop below 42.3 % for 9 mW/cm^2^, 24.7 % for 18 mW/cm^2^ and 25.7 % for 30 mW/cm^2^. During rediffusion, the concentration reaches 46.1 % for 9 mW/cm^2^, 34.9 % for 18 mW/cm^2^ and 33.9 % for 30 mW/cm^2^. On the other hand, for 300 μm the lowest values of oxygen during depletion are comparable to these of 200 μm, showing significant improvement. During rediffusion, the concentration reaches 47.5 % for 9 mW/cm^2^, 23.48 % for 18 mW/cm^2^ and 18.9 % for 30 mW/cm^2^.

An interesting issue arises from the use of asymmetrical pulsing. In both atmospheric and hyperoxic environments, the minimum oxygen concentrations for 30 mW/cm^2^ in 200 μm corneal depth are slightly higher than the ones for 18 mW/cm^2^. For 300 μm this is noted only in the atmospheric environment. A suspected reason for these values might be the different “On” times used for the different irradiances. Moreover, for the 300 μm, the finding could be explained by the sensor’s threshold and the possibility of less accurate detection since it is very close to it.

Based on the different examined protocols, pulsing seems to be efficient only for hyperoxic environment. During the symmetrical 10’’:10’’ protocol, the 9 mW/cm^2^ irradiance seems to be the optimal choice. For the asymmetrical pulsing, the irradiance of 9 and 30 mW/cm^2^ exhibit the highest oxygen concentrations.

Despite these results, this study exhibits several limitations. The first one is the *ex vivo* nature of the experiments with porcine eyes, which may not resemble completely the physiological conditions within the cornea. Additionally, the sensitivity of the oxygen micro-sensor when measuring concentrations below 1%, limits conclusions for subtle oxygen changes. Moreover, there is a bigger variability of minimum and maximum period values for oxygen measurements in the hyperoxic environment which has been previously described^17^. Another limitation of the study is the absence of biomechanical tests to determine whether and how the different pulsing protocols affect corneal biomechanics. Moreover, the treatment times for asymmetrical p-CXL could potentially be a limitation for surgeons, however, the oxygen concentration availability could compensate for the treatment time. Finally, the numerical algorithm used here is unable to characterize the oxygen consumption/performance of continuous irradiance CXL. Therefore, the comparison between asymmetrical p-CXL and continuous CXL is based on two different computational algorithm. The suggested values by the algorithm are purely based on oxygen availability and time and do not include the formation of cross-links or the saturation of cross links as a parameter and should be included in the future. For the hyperoxic situation, the algorithm must be further calibrated to suggest optimal values.

In conclusion, based on the different examined protocols, pulsing is efficient only for hyperoxic environment. During the symmetrical 10’’:10’’ protocol, the 9 mW/cm^2^ irradiance appears to be the optimal choice. For the asymmetrical pulsing, the irradiance of 9 and 30 mW/cm^2^ exhibit the highest oxygen concentrations. The results presented here provide new information on the oxygen kinetics during riboflavin-mediated symmetrical and asymmetrical p-CXL.

## Acknowledgements

We thank Avedro Inc. and Glaukos Corp. (Waltham, MA, USA) for the material support with Vibex rapid, Peschke Trade (Huenenberg, Switzerland) for the allocation of the PXL UV-source and Ziemer Ophthalmic Systems (Port, Switzerland) for their support with the channel creation using the Z8 laser system.

